# Cognitive ethology of nest building in a shell-dwelling cichlid

**DOI:** 10.1101/2025.07.27.659443

**Authors:** Swantje Grätsch, Alessandro Dorigo, Vaishnavi Agarwal, Ash V. Parker, Manuel Stemmer, Isabela Hernández Murcia, Abdelrahman Adel, Alex Jordan, Herwig Baier

**Affiliations:** Max Planck Institute for Biological Intelligence, Martinsried, Germany; Indian Institute of Science Education and Research, Bhopal, India; Bernstein Center for Computational Neuroscience, Berlin, Germany; Max Planck Institute for Animal Behavior, Konstanz, Germany

## Abstract

Across animal taxa, nest-building behavior is performed using a generalizable and flexible mental representation of an action sequence – a ‘schema’. In order to accomplish its goal (a stable nest), the brain appears to compare intermediate steps in the process to stored mental images, or ‘templates’. Deviations from these templates drive progress, preferences, and corrections in the execution of this behavior. The stereotypy vs. plasticity of both schema and mental templates have rarely been defined in an experimentally accessible system. Here we investigated nest building by the cichlid *Lamprologus ocellatus*, a fish species that manipulates abandoned snail shells to build shelters for breeding and protection. We find that nest building is composed of a sequence of behaviors that are tied together by a series of stimulus-response loops, allowing for restarts and shortcuts as the behavioral program unfolds. The attraction to a shell object is innate, as is the final appearance of the nest. The behavior in an inexperienced animal is initially uncoordinated, but is fine-tuned by repeated building opportunities. Shells need to conform to rigid geometric criteria in order to be acceptable as a potential home. Nest building is accompanied by focused neural activity in brain regions homologous to the mammalian hippocampus and neocortex. In conclusion, we have uncovered the constraints and flexibility of cognitive template matching underlying an instinctive, goal-directed behavior.

## INTRODUCTION

Nest-building behavior has evolved in a diversity of taxa – from termites to spiders and from birds to the great apes^1–4^. Depending on ecological needs and affordances, as well as the animals’ physical and cognitive abilities, the building process and nest architecture vary greatly in complexity (for a collection of fascinating photographs, see^5^). Some species construct their nests within the substrate, e.g., Australian rainforest frogs^6^ and green turtles^7^, or utilize gathered plant and animal matter, e.g., beavers^8^, harvest mice^9^, long-tailed tits^10^, orangutans^11^, platypus^12^, red squirrels^13^, and weaverbirds^14^. Other species exclusively use or integrate endogenous nesting materials e.g., orb-weaver spiders^15^, paper wasps^16^, swiftlets^17^, and tungara frogs^18^. Despite the absence of limbs or beaks, fish also exhibit remarkable abilities in nest construction^19–21^. Male white-spotted pufferfish modify the sandy bottom to create large circular nests with radial valleys and rings using their bodies^22–24^. Other species use nesting materials such as plants, algae, or animal matter to build or stabilize their nests. For example, sand gobies dig burrows under bivalve shells and add sand or mud on top of the structure using their fins or tail^25^. Additionally, some teleosts produce their own nesting materials, such as armed catfish that construct nests of mucus-covered bubbles and algae^26^, or the three-spined sticklebacks that produce the sticky ‘spiggin’ used to bind algae for their nests^27–29^.

Even though nest-building behavior has been studied for decades, we are only beginning to understand underlying neural and cognitive functions that are involved. Historically, nest building has been described as a strictly innate behavior, with little consideration given to the involvement of cognitive processes such as learned preferences, predictions, or memory^10,30^. While this theory may hold for some species, an increasing body of research suggests that cognitive plasticity plays a significant role, especially in avian nest-building behavior^2^. Many species adjust their nests to external factors, such as temperature, oxygen levels, or predation risk, indicating flexibility in the building process. Moreover, weaverbirds show various improvements with subsequent building opportunities: dexterity, morphometric variation, and material usage all improve with experience^31,32^. Studies also showed that zebra finches adapt their nesting materials based on the success of previous nests^33^ and repeated nest building of experienced zebra finches showed lower morphometric variation and usage of materials than inexperienced animals^34^. These findings collectively indicate that learning and planning are integral in nest-building and that this sophisticated behavior invites a merger of ethological concepts with those developed in cognitive science.

Here we explored plasticity and stereotypy of nest-building behavior in the cichlid fish *Lamprologus ocellatus*, which is endemic to Lake Tanganyika^35–37^. *L. ocellatus* are obligate shell-dwellers, exclusively breeding in empty snail shells and caring for their developing offspring^38^. Both males and females build nests by manipulating empty snail shells, inserting them apex-down into the substrate, and covering them with sand^36,39,40^. We discovered that the nest-building sequence of *L. ocellatus* is malleable to different shell geometry (e.g., sinistral shells), although clear innate preferences, or cognitive templates, exist. The behavioral sequence is governed by stimulus-response loops and fine-tuned by experience, suggesting a mental representation of the goal and a plan to achieve it. When performing immediate-early gene (IEG) mapping via third-generation in situ hybridization chain reaction (HCR) in the intact adult brain, we localize neural clusters active during nest building in brain regions homologous to the mammalian hippocampus and neocortex.

## RESULTS

### *L. ocellatus* builds its nest by performing a stereotyped sequence of behaviors

Nest-building in *L. ocellatus* follows a stereotyped sequence of behaviors^40^ (Figure 1E, Video 1), where fish position an empty snail shell apex-down into the substrate, cover it with sand, and keep the aperture accessible (Figure 1A, Video 2). We recorded the nest-building sequence (Figure 1D) of adult females (n = 7; Figure S1A) with a 3D-printed shell^41^ (Figure S1B) in isolated conditions (Figure 1A). Analyzing the sequence through computer vision, object detection, and manual scoring, we defined four building phases: Initiation (yellow), positioning (blue), covering (purple), and maintenance (pink) phase (Figure 1Di). In a new environment without hideouts, *L. ocellatus* often dive headfirst into the sand (in 6 of 7 experiments), remaining hidden for several minutes (‘Hiding’ in Figure 1E). Once emerged, they enter the shell within seconds (6.6 ± 3.4 s; mean ± SD) and stay close to it throughout the building sequence (Figure 1B; Diii). The first shell entry (engagement) marks the start of the initiation phase (Figure 1Di, yellow bar), during which the fish interact with the shell, *i.e.*, by scanning its surface, removing sand from its interior (‘Removing sand’ in Figure 1E), or moving it (‘Moving shell’ in Figure 1E). Substrate interactions (in 4 of 7 experiments) include mouth digging, where sand is picked up with the mouth and released in the periphery (‘Digging mouth’ in Figure 1E) and plowing, where fish perform undulatory body movements that propel sand backward (‘Plowing’, Figure 1E). Intermittently, the fish pause in front of the shell aperture with the pectoral and anal fins supporting the body on the substrate (‘Pausing’ in Figure 1E). Eventually, the fish spin the shell horizontally with the outer lip of the aperture lying flat on the substrate, a requirement for the transition into the positioning phase. On average, the initiation phase lasts 37.1 ± 36.7 minutes (mean ± SD; Figure 1F). The positioning phase (Figure 1Di, blue bar) begins with the first digging event (mouth or body digging) that occurs after the shell has been successfully oriented into the horizontal position. During this phase, the fish excavate a pit and position the shell apex-down into the sand. They systematically remove sand near the junction of the body whorl and the outer lip of the aperture (Figure S1C) through highly repetitive mouth digging, reaching up to 14 events per minute. During body digging, the fish propel sand towards the periphery as they move toward the shell (‘Digging body’ in Figure 1E). Once a pit is formed, the fish maneuver the shell into it by grabbing its inner lip and pushing the body into the ground, rotating the shell clockwise into place. This process repeats until the shell is positioned apex-down and the outer lip is level with the ground. Most shell movements occur during this phase and the last shell rotation defines the end of this building phase. On average, the positioning phase lasts 78.9 ± 31.9 minutes (mean ± SD; Figure 1F). During the covering phase (Figure 1Di, purple bar), the fish stabilize the shell and cover it with sand, primarily by plowing through the sand. They move away from the shell in a radial pattern, propelling sand towards it and thus achieve an even coverage (Figure S1D). This activity is less frequent than digging, with up to 6 events per minute. The visibility of the shell decreases as it is gradually covered with sand (Figure 1Dii). The covering phase ends when ≥ 85 % of the shell is covered and lasts on average 49.1 ± 10.8 minutes (mean ± SD; Figure 1F). The finalized covering of the shell marks the end of the active nest-building (total duration: 165.8 ± 59.1 minutes; mean ± SD; Figure 1F) and the beginning of the maintenance phase (Figure 1Di, pink bar) in which the fish primarily pause in front of the shell aperture, occasionally cover it, or remove sand from the interior.

**FIGURE 1.**
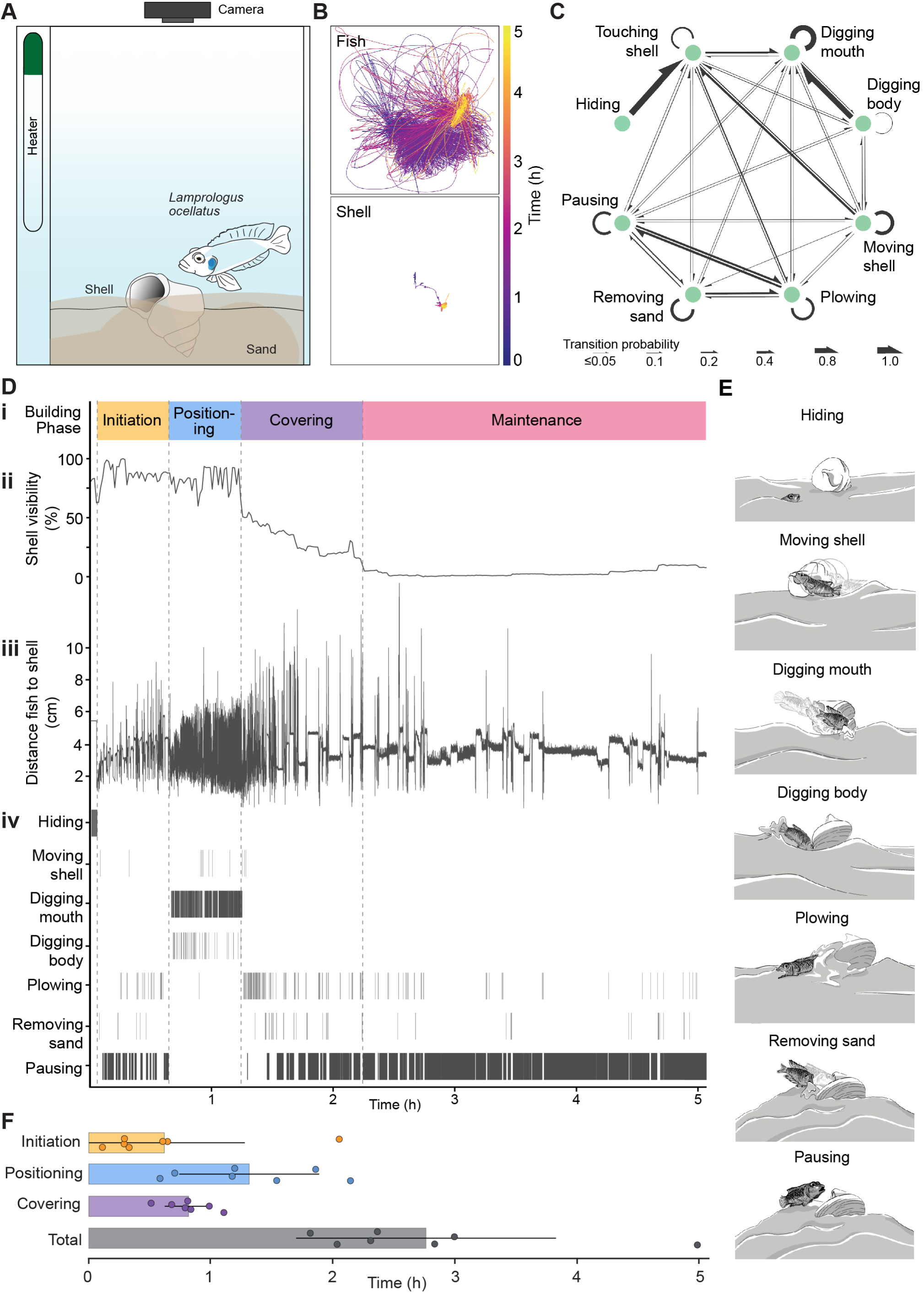
Nest-building sequence ofL *ocellatus*. A. Schematic side view of the behavioral setup showing the individual fish, a shell and the overhead camera that was used for recording the nest-building sequence. B. Trajectories of the fish (top) and the shell (bottom) during the nest-building sequence, shown over the same time course. C. Transition map of behaviors throughout the nest-building sequence. D. Summary of the nest-building sequence, including: (i) building phases; (ii) shell visibility; (iii) fish-to-shell distance plot; and (iv) behavioral ethogram over the same time course. E. Schematic representation of the specific behaviors. F. Average duration (in minutes) of the four building phases: initiation (yellow), positioning (blue), covering (purple), and total building time (grey). Values are shown as mean± SD.

In conclusion*, L. ocellatus* nest building is a robust, sequential behavior in which each construction step is closely tied to the shell’s position and orientation.

### Nest building is governed by stimulus-response loops

By externally interfering with the building phases, we tested if nest building is governed by a rigid fixed action pattern or by stimulus-response feedback loops. With the first manipulation, we aimed to reset the positioning phase by filling in the pit, excavated by the fish over the preceding 10 minutes, using sand from the surrounding area. If the animal continued digging after this manipulation, the pit was filled again after 10 minutes and this procedure was repeated up to 10 times per fish (n = 4; Figure 2B). This manipulation prolonged the positioning phase (184.4 ± 39.1 min, mean ± SD) compared to control experiments (91.6 ± 34.7 min, mean ± SD; Figure 2A and 2F), while the duration of the initiation (10.1 ± 4.6 min) and covering phases (33.8 ± 14.5 min) remained similar to control condition (11.3 ± 6.1 min and 40.9 ± 8.1 min, respectively; Figure 2F). Thus, the total building time increased to 228.5 ± 26.8 minutes (control: 144.3 ± 44.9 min, mean ± SD; Figure 2F). Notably, the fish trajectories reveal that the fish resumed digging at the same spot where they had excavated the pit before the experimental interference (Figure 2Di-v). Next, we aimed to advance the building progress and placed the shell apex-down into the sand 10 minutes after the first digging event (Figure 2C). This reduced the duration of the positioning phase to 9.9 ± 0.6 minutes (Figure 2F) and triggered an earlier transition to the covering phase (Figure 2Evii), which lasted 41 ± 21.8 minutes (Figure 2F). In total, the building time was shortened to 85.1 ± 49.2 minutes. Lastly, the shell was removed 10 minutes after the first digging event by pulling a string attached to it (Figure S2A). This caused the fish to abort digging and stay near the shell’s previous location or to hide in the sand. Animals that stayed close to the shell location performed a mixture of behaviors (body digging, mouth digging and pausing) before moving to the periphery and staying near the wall for the rest of the experiment (Figure S2A, B).

**FIGURE 2.**
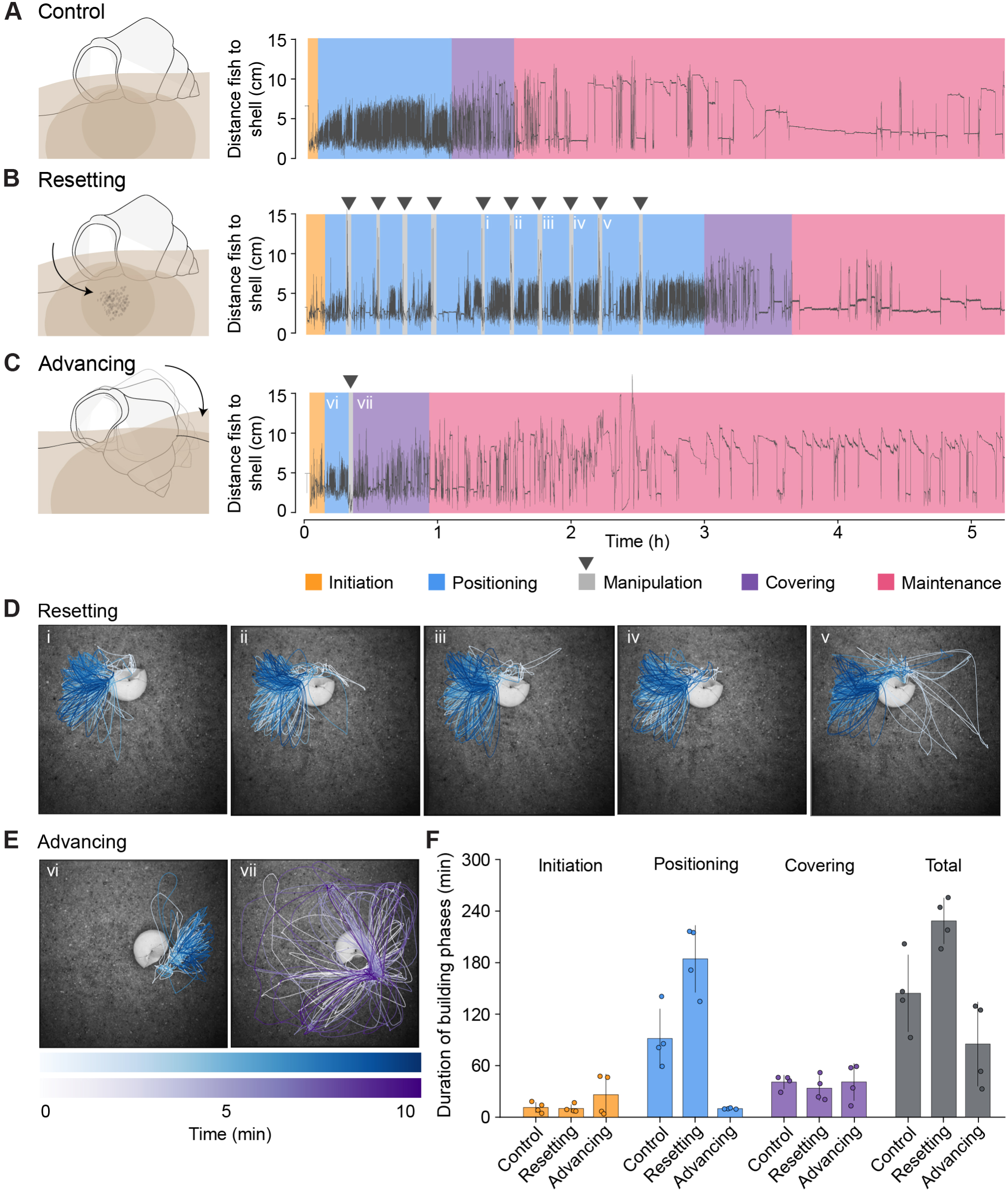
Experimental manipulations of nest-building sequence. A. Control: Fish-to-shell distance plot and the corresponding building phases during the control condition, in which the fish completed the positioning phase (blue) without interference. B. Resetting: Same representation as in A. After 10 minutes of digging activity, the pit dug during the position­ ing phase was repeatedly leveled with sand (10 events; grey shading and black arrowheads). C. Advancing: Same representation as in A. After 10 minutes of digging activity, the shell was experimentally positioned with its apex-down in the sand (grey shading and black arrowhead). D. Trajectories of the fish during the resetting condition superimposed on a photograph of the actual shell position. Panels i-v correspond to the activity marked in B. After the experimental manipulations, the fish resumed digging behavior. E. Trajectories of the fish during the advancing condition superimposed on a photograph of the actual shell position. Panels i and ii show the fish’s activity following the manipulation described in C. After the shell was placed apex-down into the sand, the fish covered it. F. Average duration (in minutes) of the building phases: initiation (yellow), positioning (blue), covering (purple), and total (grey) across control, resetting, and advancing conditions. Values are presented as mean ± SD.

Together, these findings show that fish constantly monitor shell orientation and behavioral progress, indicating that nest-building is not guided by a fixed action pattern, but by phase-specific stimulus-response-feedback loops.

### Nest building has both innate and learned components

We asked to what extent the nest-building sequence is innate. We used ‘shell-naïve’ females (n=5; Figure 3Aii) which in contrast to experienced control animals (Figure 3Ai) were reared in shell-deprived conditions. As adults, these shell-naïve females were introduced into the behavioral arena and filmed for three consecutive days. All shell-naïve fish built nests with the characteristic phenotype (Figure S3, Session 1) and executed the building sequence as described for experienced animals. However, they took longer than experienced animals to engage with the shell and complete the building sequence (Figure 3B, C). When exposed to the shell for the first time, shell-naïve animals took from 72.0 up to 1648.6 minutes to enter and engage with the shell (1006.1 ± 673.5 min; control: 0.01 ± 0.04 min, mean ± SD; Figure 3B). Before that, the shell-naïve animals hid in the sand, explored the tank, or built pits in the corner of the tank. Occasionally, they approached or touched the shell surface, but did not enter it. Once they engaged with the shell, shell-naïve fish started the nest-building sequence but took longer to complete the first stable nest (total building time: 748.2 ± 523.7 min) than experienced fish (167.8 ± 68.8 min, both mean ± SD; Figure 3C). In the positioning phase, the shell-naïve animals excavated the depression at the junction of the outer lip and the whorl, as observed for experienced animals. However, they handled the shell with less dexterity and attacked it at various places, while experienced fish mostly performed rotation movements by attacking the inner lip. Finally, 4 out of 5 nests resembled those of experienced fish (apex-down and covered). In the remaining case, the shell was positioned apex-down but was not fully covered.

**FIGURE 3.**
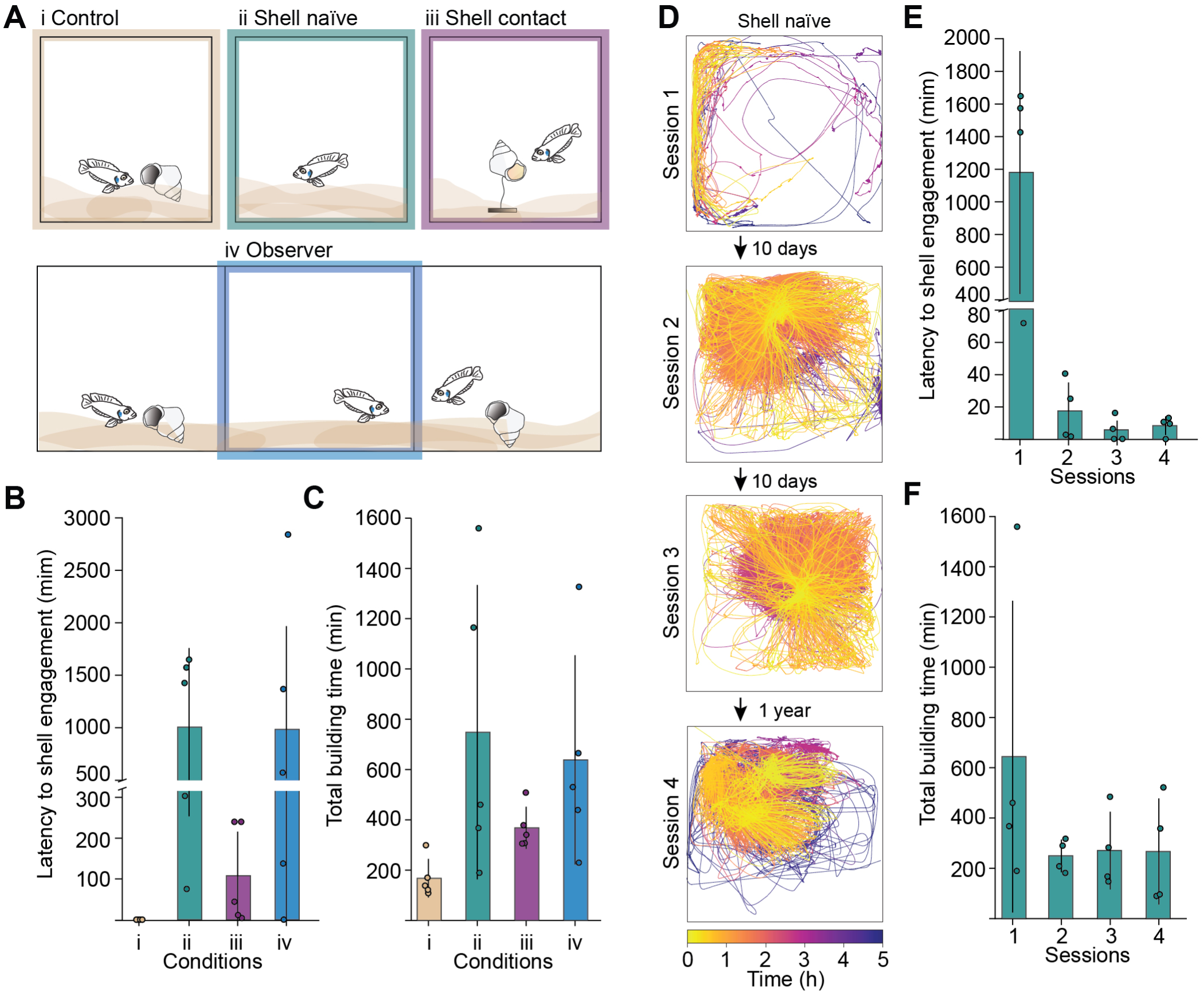
Innate and learnt components ofL *ocellatus* nest building. A. Schematic representation of the different rearing conditions. Brown: control animals; Teal: shell-naïve animals raised in a shell-deprived environment; Purple: animals raised with physical contact to a shell; Blue: animals raised observing other individuals that interacted with shells. B. Summary of latency of shell engagement and C. total building time for the individual conditions. Bars represent mean and SD, circles individual animals. D. Fish trajectories of nest-building sequence for repetitive building opportunities for shell-naïve animals (Session 1 - 4). E. Summary of latency of shell engagement and F. total building time for the individual conditions. Bars represent mean and SD, circles individual animals.

Next, we gave shell-naïve animals an opportunity to physically interact with a shell, or exposed them to the sight of shells, while not allowing them to build a nest with these shells (Figure 3Aiii and 3Aiv respectively; n=5 for each condition). Fish that observed nest building of experienced animals through a transparent divider, did not show improvements in engagement time (984 ± 1044 min, mean ± SD; Figure 3B) or total building time (638.3 ± 372.2 min, mean ± SD; Figure 3C). On the other hand, fish that were in physical contact with shells, but were not able to enter or build with them, engaged earlier than shell-naïve animals (108.0 ± 108.7 min, mean ± SD; Figure 3B) and took on average 368.1 ± 75 minutes to built the first stable nest (mean ± SD; Figure 3C).

Finally, we tested in 4 of the shell-naïve animals whether repeated building improved nest-building performance. Therefore, shell-naïve animals were transferred to their shell-deprived home tank after their first building session and then reintroduced into the behavioral arena 10 days later (Figure 3D). In the second session, they engaged with the shell more quickly (17.6 ± 16.3 min; Figure 3E and 3F, Session 2) and built nests faster (249.4 ± 56.1 min) than in the first session (engagement: 1180.1 ± 644.8 min; building time: 644.2 ± 537.3 min; Figure 3E and 3F, Session 1), and shell manipulations were more dexterous compared to the first session. In the third session, the latency to shell engagement decreased to 5.8 ± 6.6 minutes (Figure 3E, Session 3) and total building time lasted for 270.5 ± 133.8 min (Figure 3F, Session 3) while the nests resembled those from the second session. Notably, when these animals were again tested after they spent one year in their shell-deprived home tank, the latency of shell engagement was at 8.4 ± 5 minutes and total building time lasted 266.5 ± 182.9 minutes. These results show that while the nest-building sequence is innate, motor skills improve with experience and can be recruited even after a year without active nest building.

### *L. ocellatus* adapts nest-building sequence to external shell geometry

Next, we examined if the fish adapt the nest-building sequence to modified shell geometry (Figure 4A). We 3D-printed three artificial shell types: a shell with mirrored chirality (sinistral shell) but otherwise identical proportions as the control shell; a flat shell, where the aperture and whorl were not altered but the spire was compressed (total length: 4cm); and an elongated shell (total length: 6.5 cm). We used shell-experienced fish for this experiment. All fish (n = 4) engaged with the modified shells within 5 seconds and performed nest-building with the same behaviors as reported in control trials (Figure 4B). However, the total building time increased for modified shells (control: 114.4 ± 21.5 min; sinistral: 253.8 ± 106 min; flat: 1853.3 ± 34.6 min; long: 241.9 ± 45.4 min; mean ± SD; Figure 4C). Importantly, behavioral adjustments were made to altered shell geometries (Figure 4C). Shell handling of the sinistral shell initially involved dragging or pushing it from the apex or other parts. Over time, the fish adapted to the leftward chirality and switched to counterclockwise rotation by grabbing the inner lip (see Video S3). For all shell types, the fish dig a pit close to the body whorl-outer lip junction, indicating that the mirrored architecture of the sinistral shell was recognized. However, for the long shell the depression was not sufficiently excavated to position the elongated spire apex-down. Even though the fish performed prolonged digging activity, the shells were finally covered in a horizontal position. These results suggest that the elongated geometry presents a physical barrier that prevented the fish from positioning the shell apex-down.

**FIGURE 4.**
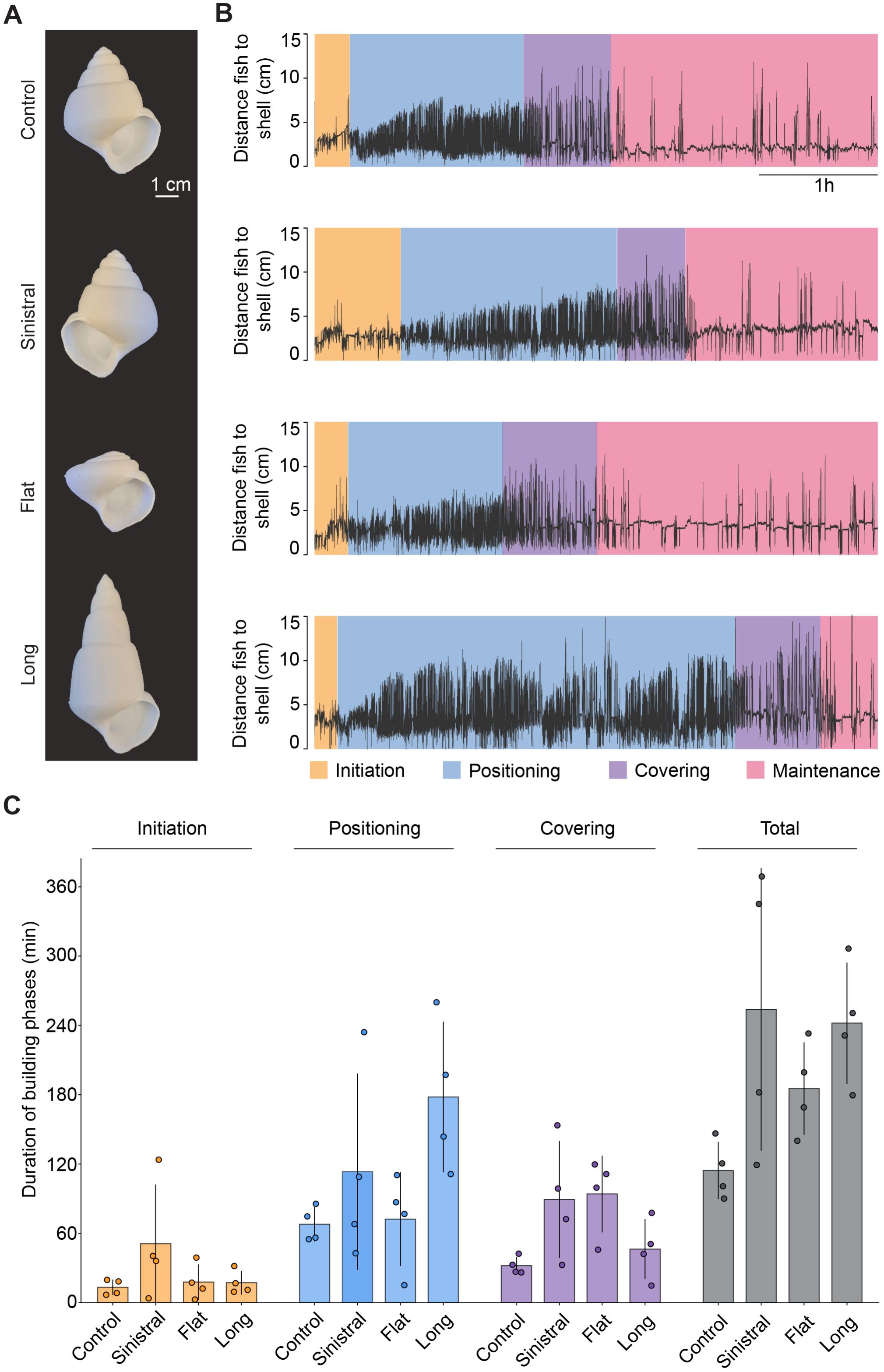
Adaptability of the nest-building sequence to different shell architectures. A. Photographs of the four shell types used, from top to bottom: control, sinistral, flat, and large. B. Corresponding nest-building sequences in response to the different shell types, represented by fish-to-shell distance plots and the associated building phases (initiation: yellow; positioning: blue; covering: purple; total building time: grey). C. Average duration (in minutes) of the building phases: initiation (yellow), positioning (blue), covering (purple), and total building time (grey) across the four shell types: control, sinistral, flat, and large shell. Values are presented as mean± SD; circles represent individual animals.

### Interior geometry and relative size are critical for *L. ocellatus* shell preference

To further explore which features make a shell suitable for *L. ocellatus* nest building, we conducted preference tests (Figure 5A) in isolated conditions using 3D-printed shells with external and internal modifications (Figure 5B). To test for size preferences, individual fish (experienced or shell-naïve females) were exposed to three shells of different length (control: 5 cm, large: 6 cm, small: 3.5 cm). Experienced females (n = 10; Figure 5C) spent most time at the control (52.9 ± 35.1 % of total time; all values in the following represent mean ± SD % of total time) and the large shell (39 ± 36 %), built them into nests, and used them for night-time resting (Figure 5D). Similarly, shell-naïve females (n = 10) spent most time at the control and large shell (49.7 ± 18.2 % and 29.8 ± 12.6 %, respectively), built nests and used them for night-time resting (Figure S5A, B). Importantly, long-term observations (20 weeks; Figure S5F) of developing *L. ocellatus* juveniles (n = 6) provided with four shell size options (3 cm, 4 cm, 5 cm, and 6 cm) revealed that, as the fish grew, they consistently selected shells that best matched their increasing body size for nest building (Figure S5G)^42^.

**FIGURE 5.**
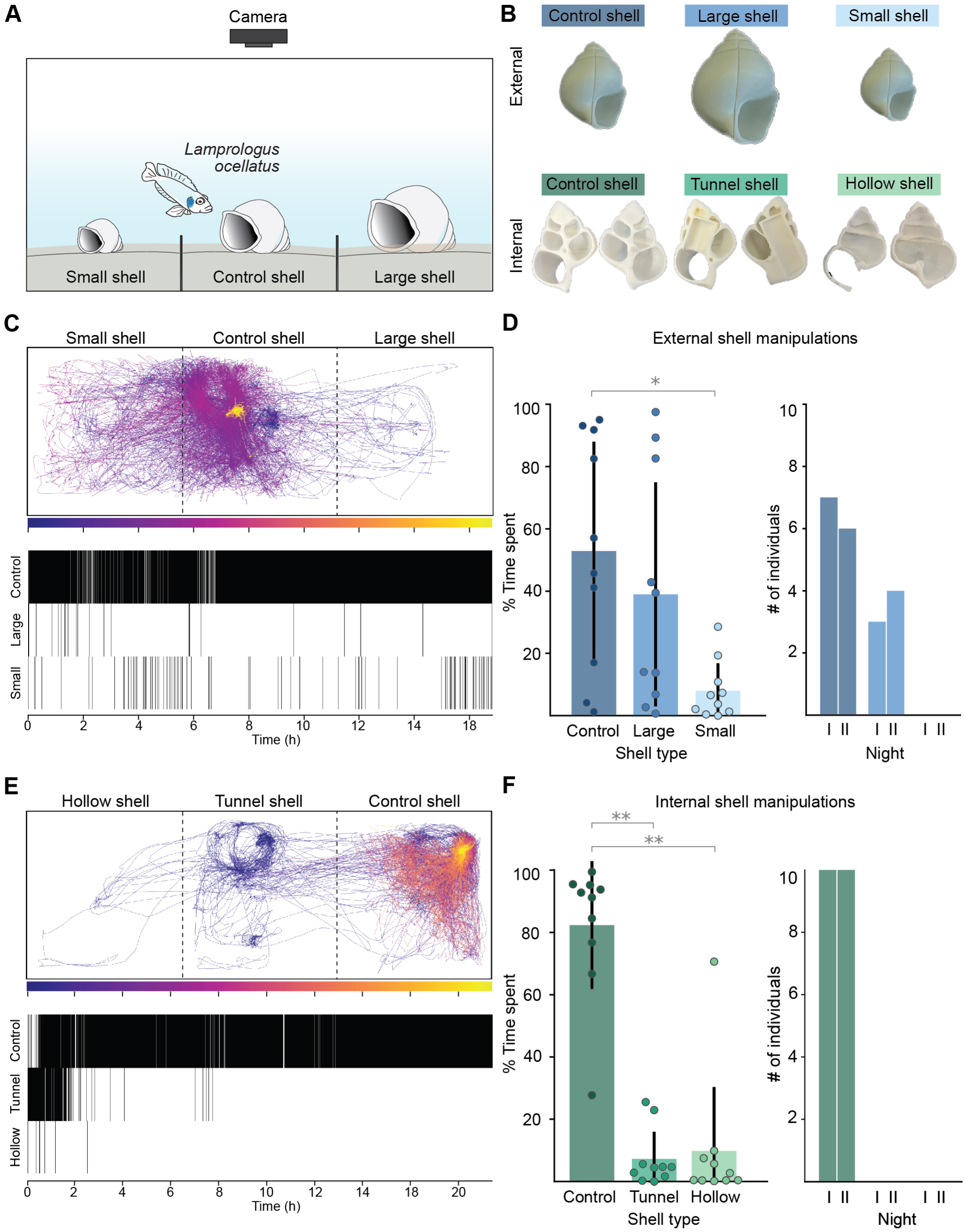
Shell preference ofL *ocellatus* for nest-building and night-time resting. A. Schematic representation of the experimental setup, showing the three individual compartments of the arena (each containing one shell), the individual fish, and the overhead camera used for recording. B. Photographs of the 3D-printed shells that were used in the experiments with manipulations of the external (top row) and internal (bottom row) architecture. C. Representative example of a fish performing preference test for externally manipulated shells. The fish trajectories (top) and corresponding ethogram (bottom) are shown, indicating the time the fish spent in each shell compartment over the same time course. D. Average time spent near each shell type during the day (left) and at night (right). Values are presented as mean ± SD. *p < 0.05. E. Representative example of a fish performing the preference test for internally manipulated shells. The fish trajectories (top) and corresponding ethogram (bottom) are shown, indicating the time the fish spent in each shell compartment over the same time course. F. Average time spent near each shell type during the day (left) and at night (right). Values are presented as mean± SD. **p < 0.01.

For internal preferences, fish were offered three shell types: control, ‘tunnel’ (columella replaced by a straight canal, and ‘hollow’ (columella removed) shell. The experienced fish (Figure 5E, F) spent most time (82.3 ± 20.5 %) at the control shell, built them into nests and used them for night-time resting. Individuals spent comparatively little time at the tunnel shell (7.3 ± 8.7 %) and hollow shell (9.8 ± 20.5%). For shell-naïve females (n = 10), we observed similar results to the control animals: the fish spent most time at the control shell (64.9 ± 19.8 %) and comparatively less time at the tunnel and hollow shell (18.4 ± 13.5 % and 16.5 ± 10.3 %, respectively; Figure S5B). Additionally, we performed the interior trial with females (n = 6) that were reared with hollow shells (Figure S5C). In housing conditions, these fish used the hollow shells and embedded them in the sand. Interestingly, these shells were not positioned apex-down but horizontally and filled with sand, reducing the internal space inside the shell. In the preference test, these females interacted more with the hollow (23.6 ± 12 %) and tunnel shell (25.3 ± 31.1 %) compared to control animals, but finally chose the control shell for nest building and night-time resting (50.8 ± 25.2 %). Overall, *L. ocellatus* selected shells appropriate to their body size, and the presence of a columella was a critical determinant of shell preference in all tested conditions.

### Dorsal telencephalic neurons are active during nest building

To investigate neural correlates of nest building, we performed in situ hybridization chain reaction (HCR) on intact adult brains (Figure 6; Video S4-6). To identify active neurons and brain structures, we examined the expression pattern of *cfos*, an immediate early gene, and *elavl3*, a marker of most differentiated neurons, using volumetric fluorescent light sheet imaging of cleared brains (Figure 6A-C). Additionally, we labeled *vglut2* and *gad2* positive cells alongside *cfos*, to identify through confocal imaging whether activated telencephalic neurons are excitatory (glutamatergic) or inhibitory (GABAergic; Figure 6D).

**FIGURE 6.**
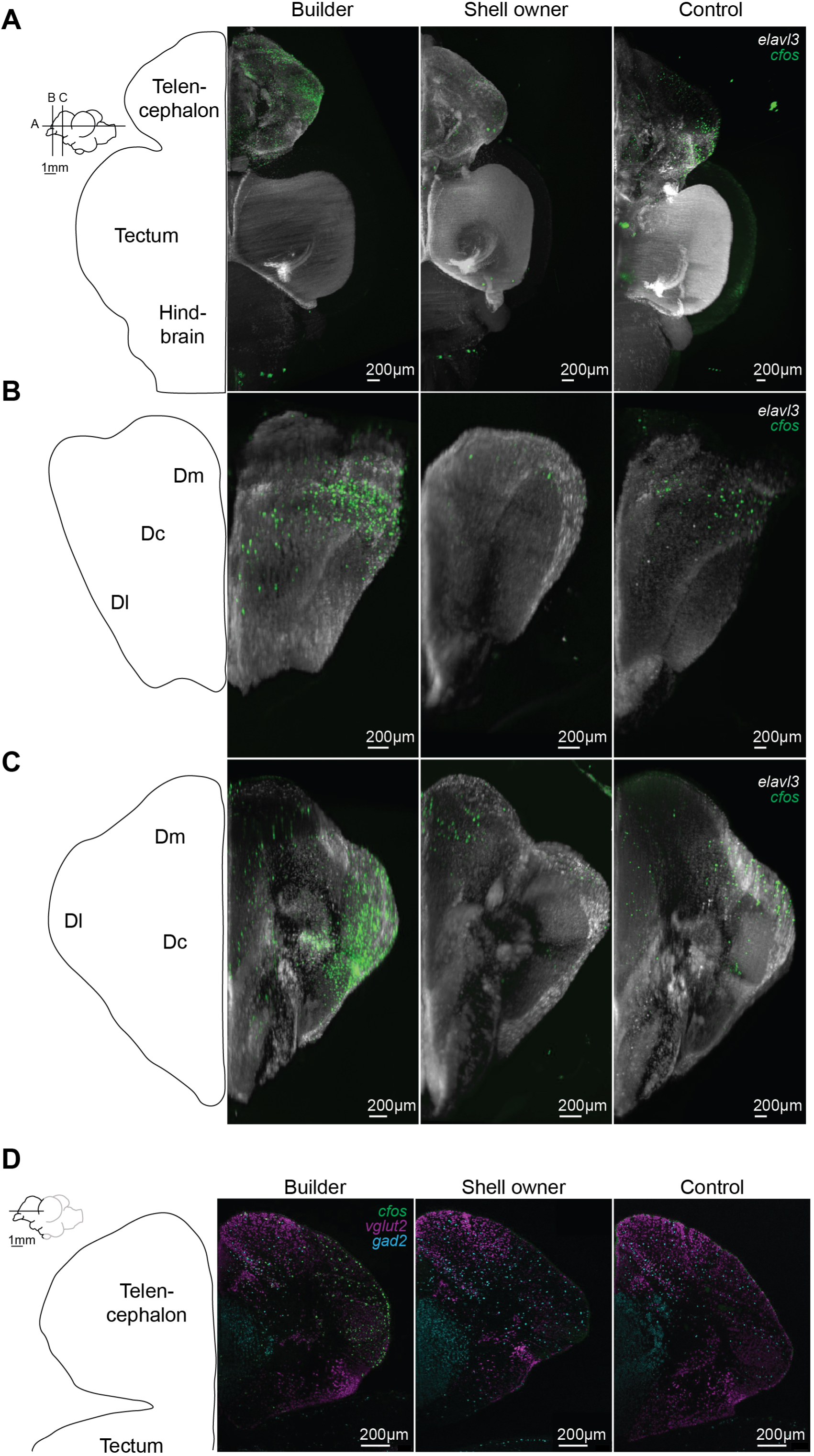
cfos-positive neurons in the telencephalon during nest-building behavior. A. Horizontal whole-brain view showing neurons positive for *e/av/3* (white), a pan-neuronal marker, and *cfos* (green), in a builder (left), shell owner (center), and control (right) animal. B. Coronal view at the level of the anterior telencephalon showing *e/av/3-* (white) and cfos-positive (green) neurons in a builder (left), shell owner (center), and control (right) animal. C. Coronal view at the level of the intermediate telencephalon showing *e/av/3-* (white) and cfos-positive (green) neurons in a builder (left), shell owner (center), and control (right) animal. D. Horizontal view of the telencephalon showing cfos-positive (green) neurons along with *vg/ut2* (magen­ ta) and *gad2* (cyan) expression in a builder (left), shell owner (center), and control (right) animal.

Each experiment included three groups of adult females (n = 4 per group): i) builders, performing nest building using a control shell, ii) shell owners, placed in an isolated tank with an experimenter-prepared nest, and iii) control animals, placed in an isolated tank without a shell but with the option to hide in the sand. After the fish in the first group engaged in digging activity for at least 45 minutes, all fish were collected from the experimental setups, euthanized, and their brain tissue was cleared and stained using the HCR protocol (see Methods). Volumetric light sheet imaging allowed us to image the complete, intact adult brain (Figure 6A-C) and observe *cfos* expression (green) in combination with pan-neuronal *elavl3* (white) expression. In nest-building animals (Video S4), we detected increased *cfos* expression in several telencephalic regions, including the dorsolateral and dorsocentral telencephalon (Dl and Dc; Figure 6B and C) and the ventral telencephalon (Figure 6A). HCR *cfos* signals were significantly less pronounced in shell owners and control animals (Video S5 and S6, respectively). Confocal images of the telencephalon (Figure 6D) revealed that *cfos*-positive neurons (Figure 6D; green) were both glutamatergic (Figure 6D; magenta) and GABAergic (Figure 6D; cyan), suggesting that multiple neuronal cell types contribute to nest building. Taken together, our whole-brain approach enabled the identification of brain regions associated with nest-building activity, highlighting regions of the telencephalon as key structures involved in this behavior.

## DISCUSSION

A growing body of evidence implicates cognitive processes, such as sensory template matching and innate or acquired schemata in nest-building activities^43^. Animals seem to “know” what nesting materials to employ –a form of tool use^44^– and what the end product should look like. Here we investigated the behavioral action patterns and decision-making in a cichlid species, *L. ocellatus*, that builds nests reproducibly in a laboratory setting. We show that nest building of experienced *L. ocellatus* animals proceeds through a stereotyped sequence of phases, which are each composed of a small number of distinct behaviors. Furthermore, we show that nest-building behavior does not represent a fixed action pattern, i. e., the unfolding of an invariant behavioral program^45^. Rather, through experimental interventions, we were able to reset, advance, or abort the nest-building sequence. Phases can be skipped or repeated, and individual behaviors are chained up in a flexible order. Together, we show that nest-building behavior in *L. ocellatus* is a concatenation of phase-specific stimulus-response feedback loops.

We asked to what extent nest building by *L. ocellatus* is innate. In birds, both instinct and learning contribute to the nest-building routine and its refinement. While inexperienced birds construct nests characteristic of their species^11–13^, the outcome of previous breeding events can influence the choice of future nesting material^33^ and nesting sites^46^. We observed that *L. ocellatus* that were reared in a shell-deprived environment instinctively used shells and built nests very similar to experienced animals. However, they engaged much later with the shell and the initial building sequence of these animals was less well choreographed and took substantially longer than in experienced animals. After a first trial, however, the nest building was fully developed and engagement latency as well as building time improved in the following trials. Observation of nest building by conspecifics without opportunity for own motor practice, did not accelerate the behavior in first-time nest builders. We conclude that nest building in *L. ocellatus* has both instinctive and learned components. Attraction to a shell, the behavioral nest-building sequence, and the final appearance of the nest are innate, while the internal organization of the behavioral program requires motor learning.

*L. ocellatus* are obligate shell-dwellers: Both in the wild and in the laboratory, they do not breed on substrates and always prefer shells over any other hollow object offered. This fixation on a shell template is species-specific: Related facultative shell users are less choosy and may breed in short pieces of tubing or in rock crevices (e. g., *Neolamprologus multifasciatus*^41^). We asked what constitutes a “shell” to the fish’s cognitive apparatus. When offered artificially altered shells, the fish readily accepted altered external geometries, including sinistral, flat or elongated shapes (although they may struggle more to maneuver the altered shells into the correct orientation). The fish also accepted larger or smaller-than-normal shells, although they preferred shells that match their own body size. However, they rejected shells in which the columella was removed or those where it was replaced with a straight tunnel. Thus, the sensory template for acceptable nesting material is an object with a helix-shaped cavity. Decisions were taken after a brief inspection and manipulation of the object on offer. It will be interesting to define the sensory modalities that enable the fish to judge and select the appropriate shell and how sensory inputs are compared to the stored template.

Unlike the learned schemata typically studied in primates and rodents^47^, the nest-building schema in *L. ocellatus* is to a large extent innate. Still, its adaptability to affordances and its flexibility in the face of perturbations suggest a cognitive contribution. In mammals, neuronal circuitry in telencephalic areas, particularly hippocampus, entorhinal, and prefrontal cortex, have been associated with geometric representations and executive control functions, including schemata of goal-directed actions. We asked if nest-building behavior was associated with activity in areas of the teleost brain that are homologous to the hippocampus and neocortex. Indeed, through whole brain mapping of previously active (*cfos*-positive) neurons, we identified neuron clusters in dorsolateral (Dl) and dorsocentral nuclei (Dc) of *L. ocellatus* that were selectively activated when the animal was building a nest, but not when it was merely maintaining it. These clusters encompassed both GABAergic and glutamatergic neurons. This result extends a previous study, which implicated glutamatergic neurons in Dl in bower-building activity of a mouthbrooding cichlid from Lake Malawi^48^. Lesion studies in cichlids (*Tilapia mossambica*), three-spined sticklebacks (*Gasterosteus aculeatus*), and Siamese fighting fish (*Betta splendens* Regan) have implicated the forebrain in the coordination of nest building^49–51^. Electrical stimulation of Dc and the ventral preoptic region elicited nest-building behavior in bluegills (*Lepomis macrochiru*s)^52^. In conclusion, these studies and our new results localize circuitry for cognitive control of nest-building activity in the teleost pallium.

## MATERIAL AND METHODS

### Husbandry and maintenance

All animal experiments were conducted in accordance with the regulations of the Max Planck Society and the regional government of Upper Bavaria (Regierung von Oberbayern), under the approved protocol ROB-55.2-2532.Vet_02-22-59. Experimental animals were bred at the Max Planck Institute for Biological Intelligence (Martinsried, Germany), from colonies established from wild-caught *Lamprologus ocellatus* collected at Isanga Bay, Zambia. For this, one adult male and two adult females were housed in 51-liter breeding tanks (600 × 270 mm, 320 mm deep). Each breeding tank was filled with 4 - 5 cm of white beach sand (grain size: 0.4 - 1.4 mm) and equipped with one 3D-printed shell (*Neothauma tanganyicense*) per fish (3D models adapted from^41^). Throughout breeding and experimentation, fish were kept under controlled conditions (13:11 h light-dark cycle; 27 °C; pH 8.2; conductivity 550 μS) and fed live Artemia daily. Room lighting was on between 7:00 am and 8:00 pm; at night, only dim light from aquarium monitors remained.

Upon reaching adulthood, experimental females (n = 53; total body length: 3.4 - 4.5 cm) were individually tagged with visible elastomer marks^53^ and transferred to 100-liter holding tanks (660 x 420 mm, 360 mm deep), where they were housed in polygynous breeding groups consisting of one male and five females. At least one 3D-printed shell was provided for each individual.

### Shell-naïve animals

Shell-naïve animals were raised in a shell-deprived environment. For this, clutches were extracted from the shells in breeding tanks before hatching (1-2 dpf), transferred into petri dishes containing facility water enriched with methylene blue (1 mL of methylene blue stock solution (0.5 g/100 mL) per 10 L of system water) and raised in incubators. The medium was changed on a daily basis and dead eggs / embryos were discarded. As soon as the larvae were swimming freely (10 dpf), they were transferred into 8 L tanks of the Tecniplast system and raised without physical or visual contact to shells, under the controlled conditions described above. For hygienic and health-monitoring reasons, no sand was present in the Tecniplast tanks. At six months old, breeding groups were selected and transferred into a shell-deprived holding tank, including sand. As they reached adulthood (one year old), they were labeled with elastomer tags and used for behavioral observations.

### Animal raised with inaccessible suspended shells, hollow shells, or visual contact to shells

Shell-naïve animals were raised as described above. After six months, breeding groups were selected and one of the following three treatments was applied for the following six months: i) the animals were exposed to silicone-sealed and apex-up suspended shells; ii) the animals were raised with hollow shells; iii) the animals were kept in holding tanks observing building animals but without access to shells.

### Designing and 3D printing of shells

3D models of the original snail shell (*Neothauma tanganyicense*)^41^ were printed (Formlabs 2 and Formlabs 3 resin printer) using white resin (Formlabs White Resin V4). Modifications of the shell architecture were performed using Blender 2.81 and Autodesk Fusion 2.0.20494 software. Unnaturalistic manipulations of the shell architecture included: sinistral shell (mirrored and thus shell of left chirality), and keeping the aperture and the body whorl unchanged, the spire of the control shell (33 mm length) was flattened for the flat shell (13 mm length) and prolonged for the long shell (53 mm length). For the preference task, three different shells were provided for each of the two trials. External manipulations implied proportional changes of shell size: control shell (50 mm), small shell (35 mm length), large shell (60 mm length); internal manipulations implied changes in the internal shell architecture: control shell (naturalistic columella), tunnel shell (columella replaced by straight tunnel of 14 mm diameter and 20 mm length), hollow shell (columella removed). Printed materials were washed and UV-cured for a minimum of 10 minutes for each procedure to eliminate any residual toxicity prior to use with live animals.

### Filming nest-building sequence

To film the nest-building sequence under socially isolated conditions, fish were placed in an experimental tank (250 × 250 mm; 30 mm depth) that was filled with fresh facility water before each trial. To prevent visual disturbance and reflections from the heater (Sera, 50W; keeping the water temperature at 27 °C), an opaque acrylic insert (200 × 200 mm; 370 mm depth) was placed inside the tank and served as the behavioral arena. Holes in the acrylic inset (10 mm diameter) allowed water exchange between the outer and inner sides of the arena. The bottom was filled with a 100 mm layer of sand, which was leveled using a small rake prior to the start of each experiment. Uniform constant white light was provided by LED strips (Multibar 35 LED strip, Lumitronix). Fish behavior was recorded from above at 30 frames per second (fps) at 1600 x 1600 px resolution using a Point Grey Grasshopper3 camera (GS3-U3-41C6NIR) equipped with a 12 mm lens (Navitar, 12-1248), and image acquisition was managed using FlyCapture2 (v2.13.3.61). Video files were post-processed using VirtualDub (v1.10.4). The fish were transferred from their holding tank into the behavioral arena, and after a habituation phase of 15 minutes, a shell was introduced into the tank with the aperture facing up. If the fish hid in the sand after the introduction, the shell was directly placed into the tank and the video recording was started. Each trial was recorded for 8 hours, then the fish was fed and left in the tank overnight. The next morning, a picture was taken of the nest before the fish was reintroduced into its holding tank.

As shell-naïve and observing animals required more time to engage with the shell and complete the nest-building sequence, the recording duration was extended to three consecutive days. During this period, fish received fresh facility water daily and were fed according to their regular schedule. Light cycle remained identical to the conditions in the housing facility.

### Filming shell preference

Shell preference experiments were performed in a 93.8-liter tank (790 x 270 mm; 440 mm depth) with an inset (600 x 200 mm; 445 mm depth) made of opaque acrylic, serving as the behavioral arena. The inset was separated into three 200 x 200 mm compartments with barriers that were reaching out of the sand up to 30 mm. This allowed the fish to visit all compartments but prevented the shells from being moved into another compartment. The arena was filled with facility sand (up to 70 mm) and a pump system (Oase BioMaster 250) was installed outside the inset, to maintain the water quality and temperature without creating visual or water disturbance in the field of view of the camera. Four white LED light strips (LumiFlex Economy LED Leiste; LumiTronix), covered with diffusion sheets and mounted on an aluminum frame above the arena, provided daytime illumination. Infrared LEDs (LED Flex Strip IR; Synergy21) were installed to record the resting site at night. The tank was illuminated with white light from 7:00 am to 8:00 pm and the infrared lights were switched on from 8:00 pm to 1:00 am. Image acquisition was performed using a ThorLabs camera (iDS UI-3240CP-NIR-GL-T) equipped with a 6 mm lens (NMV-6WA; Navitar), recording at 25 fps. The system was operated using the uEye Cockpit application (version 4.96.0650) in combination with BONSAI (version 2.8.3).

Three shells of different designs were provided for each trial; one shell per compartment. In the first experimental run, shells of different external proportions were provided, while the internal architecture was varied in the second experimental run. The positions of the shells in the tank were randomized and prepared before the fish got introduced into the arena. Control (n = 10 females) and shell-naïve (n = 10 females) fish were used for both decision tasks, fish that were reared with hollow shells (n = 6 females) were only tested for preference of the internal architecture. As one fish was introduced into the behavioral arena, the recording was started for two consecutive days for fish with prior shell experience (control and hollow) and four consecutive days for shell-naïve animals. Additionally, a photograph was taken at night to determine which shell was used for night-time resting.

### Long-term observations of shell-size preference in growing *L. ocellatus*

Two populations of juvenile *L. ocellatus* (each n = 3), aged one month, were introduced into a separate 100-liter holding tank (as described above). In each tank, 3D-printed shells (12 in total) were arranged, comprising three shells of each of four different sizes (starting shell sizes: 3 cm, 4 cm, 5 cm, and 6 cm). These shells were placed in four rows with three shells each, that were organized in a random fashion every week. The fish were allowed a one-week period to explore and select their preferred shells, during which they settled into a choice. At the end of this week, each fish was measured for body size and assigned to the shell size it occupied. The tanks were reset, including a new random organization of the shells and the fish were again given one week to choose their shells. This cycle of removal and replacement of the smallest shell size row was repeated weekly for a total of 20 weeks.

### *cfos*-based activity mapping

Adult females (n = 15) were used to map neurons active during nest-building behavior. Three animals were simultaneously introduced into separate, isolated tanks under the following experimental conditions: with a shell placed on top of the sand surface with the aperture facing up (builder), with a shell built into a nest by the experimenter (shell owner), without access to shell (control). The builder was recorded and monitored during the experiment, serving as a temporal reference: as soon as the female performed mouth digging (positioning phase), all fish were allowed to remain in their tanks for 45 minutes. The builder continued to be remotely monitored to ensure active nest-building behavior was performed throughout the entire time frame, allowing for consistent *cfos* expression. After 45 minutes, the three animals were collected, immediately euthanized and decapitated to be processed further for the in situ hybridization protocol.

### In situ hybridization chain reaction

Tissue preparation for whole-mount in situ hybridization was performed as described in the HCR RNA-FISH protocol for zebrafish embryos and larvae (Molecular Instruments). All reagents including hairpins, buffers and *cfos* probe were purchased from Molecular Instruments. *gad2* and *vglut2* probe pairs were designed using the in situ probe generator program from the Özpolat lab.

After the behavioral assay (described above), the fish were immediately euthanized and decapitated. The head was then fixed in 4% PFA/ DPBS-T (0.3% Triton X-100) for 24 hours at 4°C on a shaker. Brain dissection was performed the following day with forceps and surgical micro scissors (Fine Science Tool) in ice cold DPBS under a stereo microscope (Nikon SMZ 645) coupled with an external light source (Zeiss CL 6000 LED). Brains were then washed in 1 x DPBS-T (3 x 5 minutes), dehydrated in 100% Methanol (4 washes x 10 minutes, 1 final wash x 50 minutes) and stored at −20°C until further use. After rehydration with a series of graded Methanol/DPBS-T washes, tissue was permeabilized with Proteinase K (10 µg/mL) for 20 minutes at RT. Brains were then postfixed in 4% PFA/ DPBS-T for 20 minutes at RT. After extensive washes in DPBS-T, brains were incubated in hybridization buffer (Molecular Instruments) for 1 hour at 37°C. Subsequently, the hybridization buffer was exchanged with fresh probe solution, which was prepared by transferring 2 pmol of each HCR probe set (2 µl of 1 µM stock) to 500 µl of hybridization buffer at 37 °C. The sample was then incubated between 48 and 60 hours at 37°C on a shaker. To remove unbound probes, samples were washed in a pre-warmed wash buffer (Molecular Instruments) 4 times for 15 minutes at 37°C and finally in 5 x SSC-T (0.3% Triton X-100) at RT. Brains were incubated overnight at 4°C in a solution containing 0.2% Bisacrylamide, 3.8% Acrylamide, 0.06 M Tris-HCl, pH8. The next morning the hydrogel was allowed to polymerize with TEMED (0.2%) and APS (0.2%) in a vacuum chamber at 37°C for 2 hours. To clear light scattering lipids the specimen was incubated in a digestion buffer (2 x SCC, 2% SDS, 0.5% Triton X-100, 2mM CaCl_2_ and 1 x Proteinase K) for 24 hours at 37°C on a shaker. Samples were washed in 1 x DPBS-T and incubated in an amplification buffer for 30 minutes at RT. Finally, this was replaced by a fresh amplification buffer containing the desired fluorescent hairpins and it was further incubated for 48 hours at RT.

### Imaging of brain tissue

Images of the whole adult brain were taken using a light sheet microscope (UltraMicroscope II; LaVision BioTec) with a 4x MI PLAN objective equipped with a dipping cap for aqueous buffers. High-resolution images of the telencephalon were taken on a confocal microscope (Leica SP8) coupled with a 20x HCX APO objective and a 25x HC FLUOTAR immersion objective.

### Annotation of nest-building sequence and transition map

Video footage of the nest-building sequence was manually annotated using BORIS (Behavioral Observation Research Interactive Software^54^). The different behaviors and categorization into the building phases were adapted from previous studies in *L. ocellatus* nest building^40^. Behaviors were annotated as state events (with start and stop timestamps) and labelled as hiding, moving shell, digging mouth, digging body, plowing, removing sand, or pausing (Figure 1E, Video 1). Additionally, the building phases were annotated in BORIS and defined as initiation phase, positioning phase, covering phase, and maintenance phase.

For the transition map (Figure 1C), the number of transitions from each behavior to another were extracted for each experiment using BORIS. These matrices were then pooled and averaged using custom-written Python code. The ethogram as well as the transition map were plotted using custom-written Python codes.

### Automated tracking of fish and shell

To detect the position of the fish and the shell during nest building, we used YOLO v5^55^ and trained this object detector model using a custom-written Python GUI on 910 manually annotated images (labelImg, Github) across 27 videos, ensuring an even spread across the timeframe of each experiment. Object categories labeled with bounding boxes included one fish and one shell per image. Data augmentation was used to transform the labeled images (applying up to seven different transformations including rotations, blurring, and sharpening) to increase the size and diversity of the training data set. The model was trained to convergence and then used for inference in complete video recordings of the different experiments to predict the image position of each identified object’s center (fish and shell). Distance plots, trajectories, and bar plots were generated using custom-written Python codes.

For the decision task, the previous YOLO model was used to initialize a second round of training with 1,250 manually labeled images extracted from 25 videos. In each image, the image position of one fish and three shells were labeled. These images were further transformed through data augmentation and included in the training process. Once again, the model was trained to convergence and used for inference on the complete dataset in order to predict the image position of the fish and the three shells’ center.

### Analysis of shell visibility

To quantify the process of shell covering by the fish over the course of the experiment, we developed a custom image analysis pipeline. Video recordings were processed by extracting one frame per minute. Each frame was converted to grayscale, and the number of white and black pixels was calculated. Pixels with an intensity value of ≥ 100 were classified as white (exposed substrate), while darker pixels were considered black (covered substrate). White pixel values were normalized to the maximum value within each time series. A threshold of 15% of the maximum white pixel count was used to define the covering phase. The shell visibility was plotted using custom-written Python code.

### Analyzing time spent at a shell during preference task

To analyze the time fish spent near individual shells, three regions of interest (‘shell regions’) were manually annotated, each corresponding to a shell-assigned compartment (labelme, GitHub). These shell regions were labeled according to their relative positions in the arena, using the 20 × 20 cm compartments as visual guidance. These annotations allowed fish positions to be assigned to specific shell types over time. An analysis pipeline^38^ was adapted and used to calculate the total time each fish spent at each shell. Individual fish’s data were then grouped by experimental condition (control and shell-naïve animals) and used to compute the average percentage of time spent in each shell region. To test for shell preferences within each group, we applied two-tailed Wilcoxon signed-rank tests. A significance level of 0.05 was used for all analyses.

## DATA AVAILABILITY STATEMENT

A sample dataset is available at: https://zenodo.org/records/15593876?token=eyJhbGciOiJIUzUxMiJ9.eyJpZCI6IjcxZWUzMDE1LTFlM2ItNGIzNS05MjY4LThkMzM1MDdjMTBiOSIsImRhdGEiOnt9LCJyYW5kb20iOiI5Yjc2OTUzOGE2MDhhNzdiZWM4NDA5YjdkYWJjNDYwYSJ9.tWiwQ2fgx1HiaHh2n12gbjtqMpnzcressKNSzhR4hO9i0KF-1X_Hocuoy-6tGX9nU2suDhXXRQNGpqAXQHvEeA

Remaining video and brain imaging data sets for this study will be available upon publication.

## CODE AVAILABILITY STATEMENT

Analysis pipeline was adapted from^38^. Detailed code used for this study will be available upon publication.

## ACKNOWLEDGMENTS

The authors would like to thank current and past members of the Baier lab and colleagues at the Max Planck Institute of Biological Intelligence for collaboration, discussions, and advice at all stages of the project, especially Gregory Marquart. Kalyn Dawes devised the non-invasive shell-removal assay and collected data. Joe Donovan advised on 3D printing and data analysis. Krasimir Slanchev managed the fish facility, and his team of animal caretakers kept our cichlids healthy and well-fed. Karin Finger-Baier and Jula Huppert helped with composing animal protocols. Funding was provided by the Max Planck Society and Human Frontiers Science Program (RGP0052/2019).

## ETHICS DECLARATIONS

The authors declare no competing interests.

## AUTHOR CONTRIBUTIONS

Conceptualization, S.G., A.D., V.A., A.V.P., M.S., and H.B.; methodology, S.G., A.D., V.A., A.V.P., M.S., A.A., I.H.M. and H.B.; investigation, S.G., A.D., and V.A.; software, S.G., I.H.M., and A.A.; formal analysis, S.G., A.D., V.A., I.H.M., and A.A.; visualization, S.G., A.D., V.A., I.H.M., and A.A..; writing – original draft, S.G.; writing – review and editing, S.G., A.D., A.V.P., I.H.M., A.J., and H.B.; supervision, H.B.; funding acquisition, H.B.

## Notes

### Competing Interest Statement

The authors have declared no competing interest.

