## Supplemental Figures 1-4 for "Cognitive ethology of nest building in a shell-dwelling cichlid"

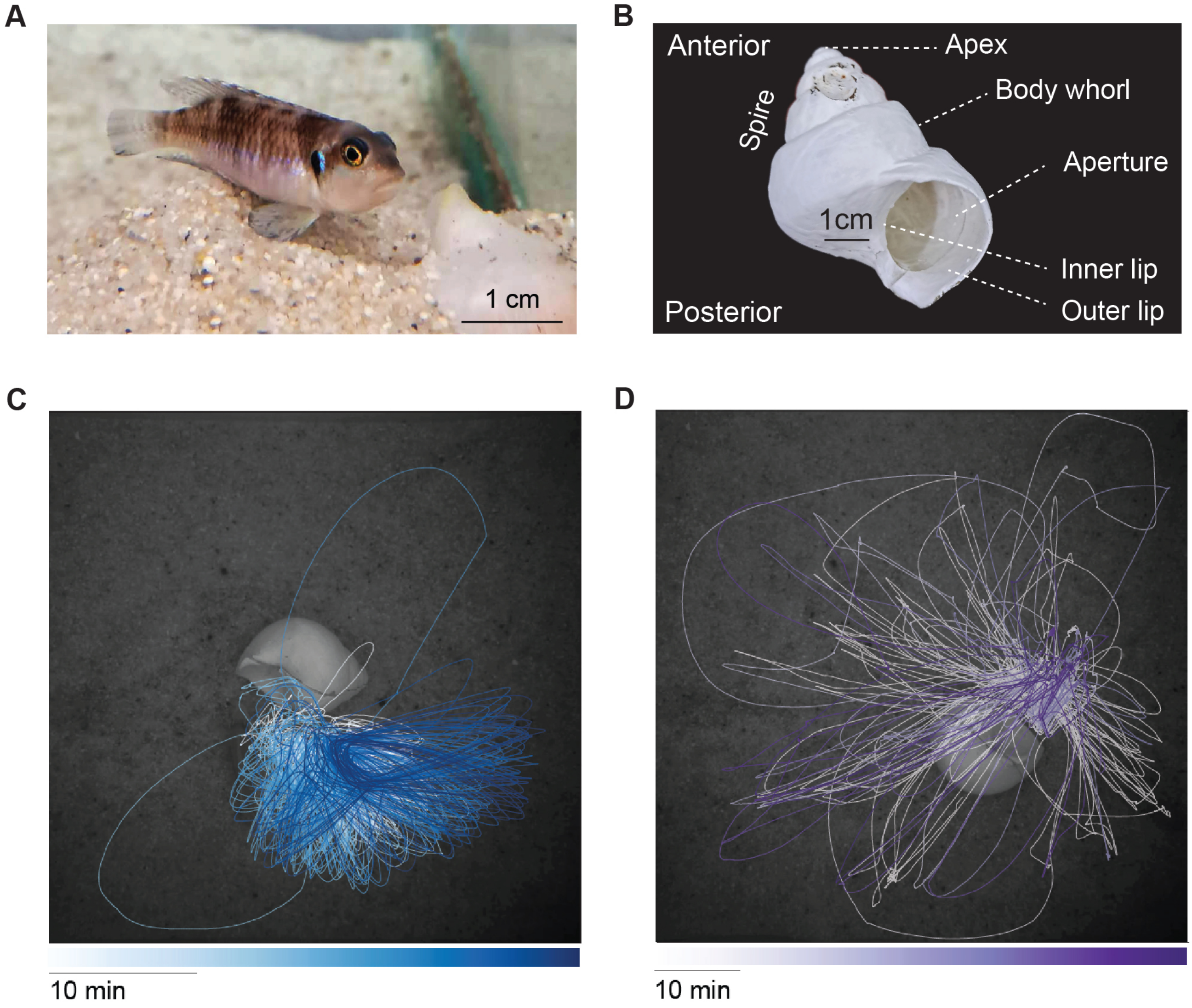


**FIGURE S1. Nest-building sequence of *L. ocellatus*.**
A. Female L. ocellatus guarding her nest.
B. 3D-printed snail shell that was used for the control experiments with dimensions and anatomical terms.
C. Trajectory of a fish's movements during the positioning phase. The trajectory of digging activity is represented as an overlay of a photograph of the actual shell position.
D. Trajectory of a fish's movements during the covering phase phase. The trajectory of digging activity is represent-ed as an overlay of a photograph of the actual shell position.

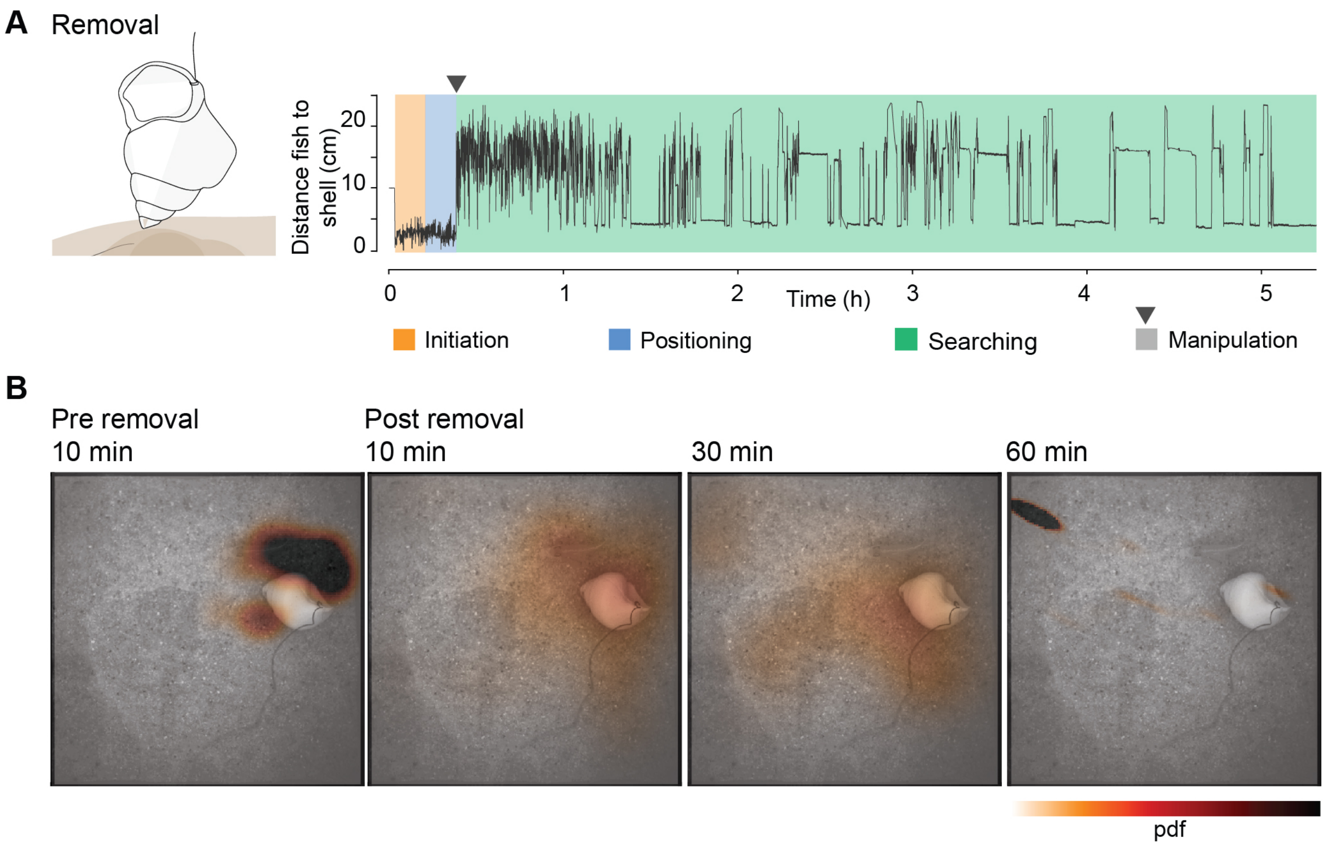


**FIGURE S2. Removal of shell during positioning phase.**
A. Removal: Same representation as in Figure 2A. After 10 minutes of digging activity, the shell was removed (grey shading and black arrowhead). After the shell was removed, the fish aborted the digging activity and searched for the shell in the vicinity (green).
B. Kernel density estimation (KDE) plot superimposed on a photograph showing the last position of the shell before removal. KDE represents location 10 minutes before and 10, 30, and 60 minutes after the shell removal. pdf: probability density function.

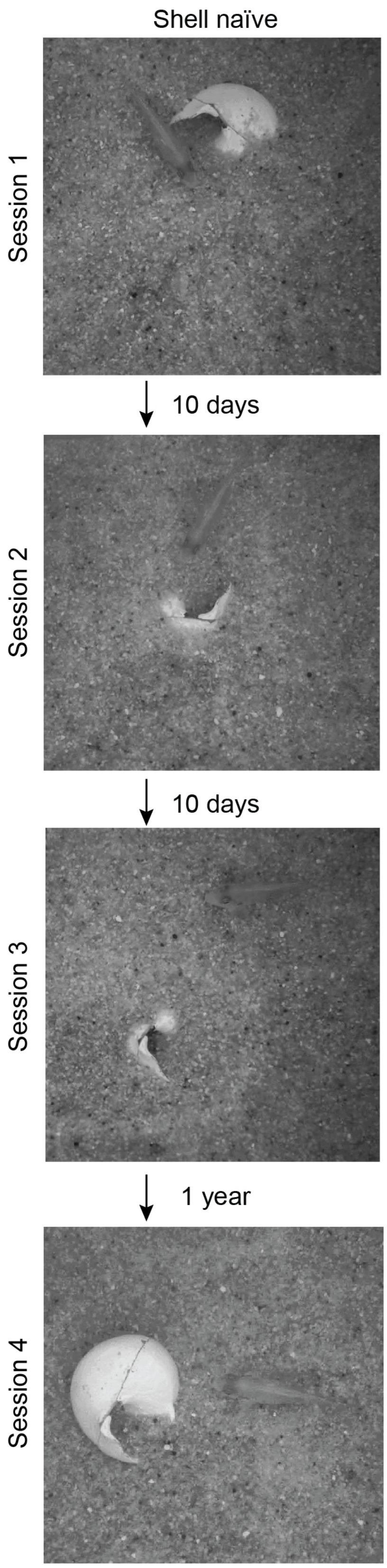


**Figure S3. Repeated nest-building behavior in a shell-naïve female.**
Photographs of the final nests built by a shell-naïve female in four separate sessions. The second and third sessions occurred 10 days after the previous one, while the fourth session took place one year after the third. Each image shows the completed nest at the end of the respective building period.

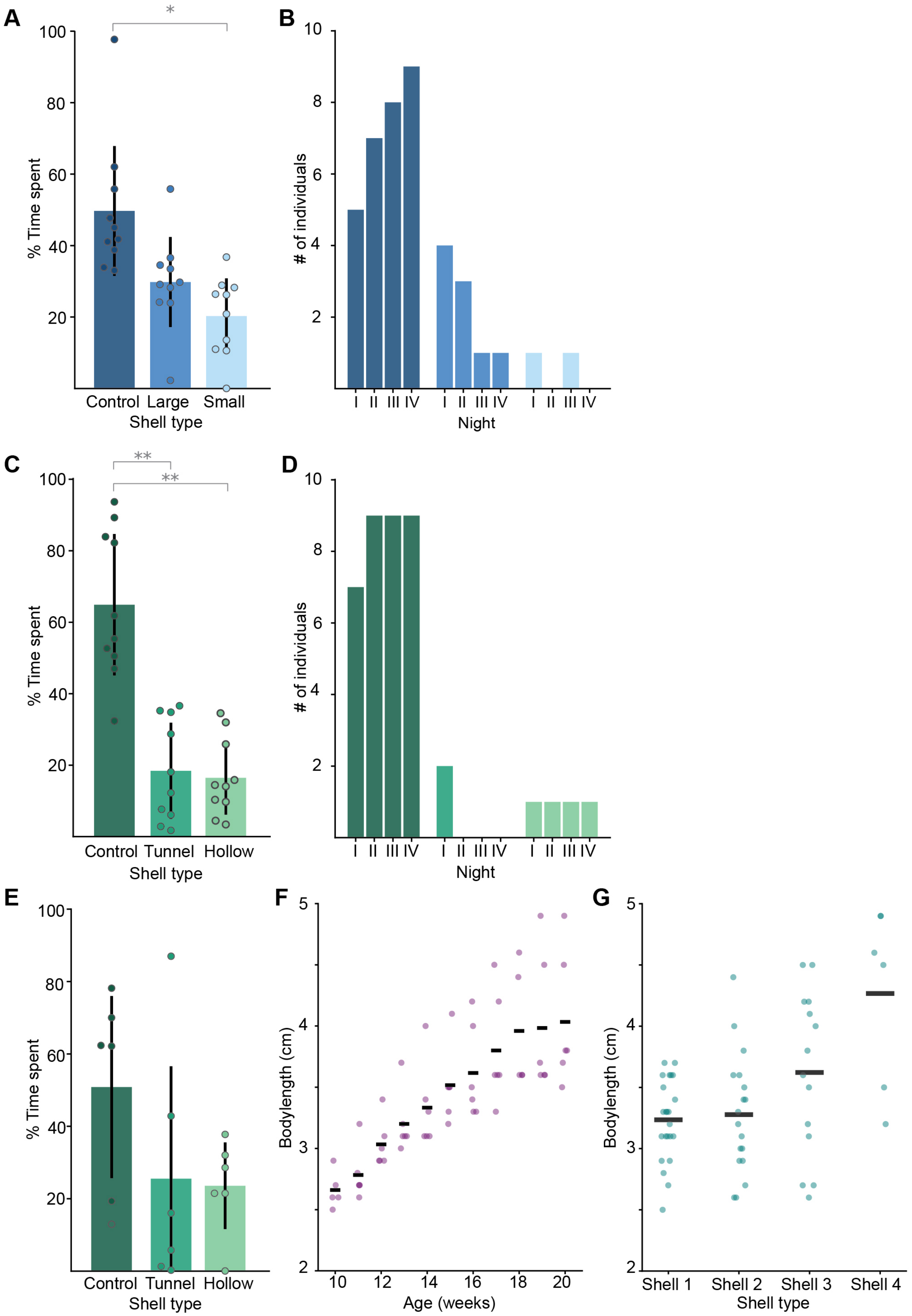


**Figure S4. Shell preference of shell-naïve and hollow-shell–reared *L. ocellatus* for nest-building and night-time resting.**
A. Average time shell-naïve animals spent near each shell type (external manipulations) during the day and B. at night. Values are presented as mean ± SD. *$p < 0.5$.
C. Average time shell-naïve animals spent near each shell type (internal manipulations) during the day and D. at night. Values are presented as mean ± SD. **$p < 0.01$.
E. Average time hollow-shell-reared animals spent near each shell type (internal manipulations) during the day. Values are presented as mean ± SD.

F. Body size of growing L. ocellatus up to the age of 20 weeks. Dots represent body size of individual animals; bars represent the mean body size.
G. Shell occupation of the individuals shown in F. Shell lengths: Shell 1 = 3 cm, Shell 2 = 4 cm, Shell 3 = 5 cm, Shell 4 = 6 cm. Dots represent body size of individual animals; bars represent the mean body size.
